# Somatotopic Organization of the Ventroposterior Nucleus of the Thalamus in Macaque Monkeys with Dorsal Spinal Cord Injuries Sustained During Early Postnatal Development

**DOI:** 10.64898/2026.10.21.759785

**Authors:** Neeraj Jain, Hui-Xin Qi, Arun Raman, David C Lyon, Jon H Kaas

## Abstract

Injuries to peripheral nerves or the spinal cord lead to changes in the topographic organization in the brain at multiple levels along the sensory pathways. In adult monkeys large-scale deafferentations due to transection of the dorsal columns of the spinal cord at cervical levels result in large-scale reorganizations such that the deafferented neurons in the hand representation of somatosensory cortical area 3b start responding to anatomically distant body parts such as the chin. However, reorganization following dorsal column injuries depends upon the age at the time of the injury. The somatotopy of areas 3b, 1 and 2 in macaque monkeys with injuries sustained on or before postnatal day (PND) 5 is very normal-like. Whereas in monkeys with injuries sustained between PND9 and PND12, a large extent of the deafferented somatosensory cortex remains unresponsive to peripheral touch, with neurons at only a few sites responding to touch on the face or arm. Here we determined the organization of the ventroposterior nucleus (VPN) of the thalamus in four of these monkeys (*Macaca radiata*, 3 females, 1 male) that received lesions of the dorsal columns on PND 3, 5 and 9. We show that the nature of reorganization of VPN largely resembles that of the somatosensory cortex. Somatotopy is normal-like after lesions on PND3 and 5, whereas after lesion on PND9, the deprived neurons are largely unresponsive to peripheral touch, with a limited expansion of the chin representation.

**Key points:** Injuries to the dorsal columns of the spinal cord in newborn macaque monkeys result in reorganization of the somatosensory ventroposterior nucleus (VPN), which depends on the age at the time of lesion. In monkeys with injuries on or before postnatal day (PND) 5, VPN is very normal-like. In monkeys with injuries on PND 9, deafferented neurons in VPN remain largely unresponsive to touch. The nature of reorganization is very similar to that previously described for the somatosensory cortex. Thus, cortical reorganization is an upstream reflection of changes in the somatosensory thalamus.

## INTRODUCTION

Information processing networks in the brain respond to changes in sensory inputs resulting from injuries. After loss of sensory inputs due to deafferentation in adult primates, topographic reorganization has been observed in the somatosensory (Florence et al., 1997), visual (Hubel et al., 1977; Kaas et al., 1990), and auditory systems (Schwaber et al., 1993). In the somatosensory system, neurons in the deafferented parts of the brainstem nuclei, the somatosensory thalamus, and the somatosensory cortex come to respond to adjacent intact sensory inputs (Dutta et al., 2014; Jain et al., 2008; Kaas et al., 1999; Kambi et al., 2014). The nature of reorganization depends upon site and extent of the injury, and age of the animal at the time of injury. For example, after injuries to a nerve to the hand in adult monkeys, the deafferented neurons in the somatosensory cortical areas 3b and 1 start responding to the intact inputs from other nerves to the hand (Merzenich et al., 1983). Deafferented parts of the somatosensory thalamus, i.e., the ventroposterior (VP) nucleus, and the cuneate nucleus of the brainstem get reactivated by same inputs that reactivate area 3b (Churchill et al., 2001; Garraghty and Kaas, 1991). After more extensive deafferentations, e.g., due to transections of the dorsal columns of the spinal cord at cervical levels, inputs from the chin, and to a smaller extent from the arm, reactivate neurons in the deafferented hand cortex (Jain et al., 1997; Jain et al., 2008). Similarly, inputs from the chin reactivate deprived neurons in VP nucleus and the cuneate nucleus (Jain et al., 2008; Kambi et al., 2014). Additionally, inputs from the occiput, neck, and shoulder reactivate deafferented neurons more extensively in the cuneate nucleus and VP nucleus, than in area 3b (Halder et al., 2018).

Reorganization in a developing brain after a sensory loss has a different outcome as compared to that in adult animals. For example, after unilateral visual deprivation early in development, during what is called the critical period, there is an irreversible change in the width of the ocular dominance columns in primary visual cortex (area 17 or area V1) (LeVay et al., 1980). In the lateral geniculate nucleus (LGN) of the thalamus, cells in the layers corresponding to the deprived eyes shrink (Sesma et al., 1984; Takahata et al., 2018).

We have recently shown (Jain et al., 2025b) that in newborn macaque monkeys, transections of the dorsal columns of the spinal cord at cervical levels result in reorganization of the somatosensory cortex which is different from that seen in adult animals after a similar injury (Jain et al., 2008). In monkeys with dorsal column lesions made on or before post-natal day 5 (PND5), somatotopy in area 3b and the adjacent somatosensory cortex, i.e., areas 1 and 2, is very normal-like. However, in animals lesioned between PND9 and PND12, a large expanse of the deafferented somatosensory cortex remains unresponsive to any peripheral stimulation, although there is a limited reactivation of the deprived neurons by inputs from the chin and the arm. Thus, cortical reorganization following dorsal column injuries early in development is different from that seen following injuries in adult animals, where neurons in the entire deprived cortex get reactivated by inputs from the chin, and to a limited extent by arm inputs in the medial cortex (Jain et al., 2008). However, it is not known how spinal cord injuries during early postnatal days affect the organization of the VP nucleus of the thalamus.

Here, we determined if alternative ascending sensory inputs reactivate neurons in the VP nucleus of the thalamus following unilateral surgical lesions of the dorsal columns of the spinal cord at cervical levels made when the monkeys were between 3 and 9 days old. Organization of the VP nucleus (ventroposterior lateral, VPL; and ventroposterior medial, VPM) was determined using standard multiunit mapping techniques when the monkeys were between 3-5 years old. Previously we have published data from these monkeys from motor cortex (Qi et al., 2010) and the somatosensory cortex (Jain et al., 2025b). An abstract of results presented here has been published in brief (Jain et al., 2025a).

## MATERIALS AND METHODS

Four bonnet macaque monkeys (*Macaca radiata*) born in the colony were used for these experiments. Dorsal columns were lesioned unilaterally at cervical levels between 3^rd^ and 9^th^ postnatal day. For details of the monkeys, see Table 1. All animal procedures were approved by the Institutional Animal Care and Use Committee of Vanderbilt University, and followed NIH guidelines. Details of the methods have been described before (Jain et al., 2025b).

**Table 1.** Details of the experimental monkeys, including the age at the time of the dorsal column lesion and when the VP nucleus was mapped, and other details. (y, years; m, months; d, days)

| Animal No. | Sex | Side of the lesion | Level of the lesion | Age at the time of the lesion | Age at the time of analysis |
| --- | --- | --- | --- | --- | --- |
| 97-63MK | Female | Left | C5 | 3 days | 3y 3m 19d |
| 98-18MK | Female | Left | C3/C4 | 3 days | 3y 11m 18d |
| 97-58MK | Female | Left | C3/C4 | 5 days | 4y 19d |
| 99-84MK | Male | Left | C3/C4 | 9 days | 4y 11m 1d |

### Dorsal column lesions

Unilateral lesions of the dorsal column were made at cervical levels of the spinal cord under sterile conditions. The animals were deeply anesthetized with a mixture of ketamine (15 mg/kg) and xylazine (0.4 mg/kg), upper back of the animals was shaved and prepared for surgery. A midline incision was made in the skin, and muscles were retracted with blunt dissection to expose the vertebral column. The level of the lesion was ascertained by counting the dorsal spines of the vertebrae. A partial laminectomy was made to expose the spinal cord, and dura was incised and reflected. The dorsal columns were transected unilaterally, from site of entry of the dorsal roots to the midline using a pair of fine forceps. Site for the lesion was selected in order to minimize damage to the surface blood vessels. For detailed procedures see Jain et al. (2008, 2025b). After lesion was made, the dura was pulled back and a small piece of gel foam was placed over the spinal cord. The muscles were retracted back and sutured in place. The skin incision was closed by intradermal sutures, and surgical glue was used for a complete closure. During the entire surgical procedure, anaesthesia was supplemented with ketamine and xylazine as needed by administering 20% of the initial dosage (IM). Core body temperature was continuously monitored and maintained with a water circulating heating pad placed under the animal. The heart rate and blood oxygenation level were monitored throughout the surgical procedure. Immediately after the surgery, animals were given dexamethasone (2 mg/kg, i.m.) and penicillin (60,000 IU/kg).

After the animals were fully awake, they were returned to their mothers for normal rearing. The monkeys were carefully observed during the recovery period. Recoveries were uneventful for all the monkeys, and mothers took care of the babies as normal.

### Multiunit mapping of the somatosensory thalamus

After approximately 3-5 years of recovery period (Table 1) somatotopic organization of the ventroposterior (VP) nucleus was determined using multiunit mapping techniques (Jain et al., 2008; Tandon et al., 2009). Monkeys were anesthetized using ketamine hydrochloride (10-20 mg/kg, IM) and xylazine (0.4 mg/kg, IM), supplemented with urethane (250mg/kg) as described before (Jain et al., 2008). The head of the animal was stabilized in a stereotaxic apparatus; the skull was exposed, and an opening was made over most of the mediolateral extent of the central sulcus. The opening in the skull was surrounded by a dam of acrylic plastic, the dura was retracted, and the exposed brain surface was covered with silicone fluid to prevent desiccation. Recordings in the VP nucleus contralateral to the lesion were made using standard multiunit electrophysiological techniques as described before (Jain et al., 2008). The procedure is briefly described below.

A photograph of the brain surface was taken and printed at high magnification on which locations of the electrode penetration sites were marked with respect to the surface blood vessels. Recordings were done using a parylene coated tungsten microelectrode (1 MΩ at 1 kHz; Microprobe, MD, USA), which was advanced down into the brain in a stereotaxically vertical plane, approximately 5 mm from the midline and 2 mm rostral to the central sulcus. Initially, the electrode was advanced using the micromanipulator while the skin all over the body was stimulated by hand to determine location of the VP nucleus. Once VP nucleus was located, this depth was noted, and the electrode was withdrawn. Subsequently, a micromanipulator was used to manually insert the electrode to a depth of about 2 mm above the plane of VPN (about 10-14 mm from the brain surface). A virtual horizontal plane at this depth was used for subsequent penetrations as a starting point for recordings. From this depth, the electrode was advanced using a hydraulic Microdrive (Model 2650, David Kopf Instruments, CA, USA), and receptive fields of the neurons were determined at 300-400 µm intervals. At each recording site, the entire body was thoroughly explored for neuronal responses to light tactile stimuli, taps, and movements of the joints and muscles. After the electrode reached a depth where neurons did not respond to tactile stimulation, it was withdrawn and inserted at a different location. The penetrations were made in a grid-like fashion, and an effort was made to cover the entire mediolateral and anteroposterior extent of the VP nucleus.

The neuronal responses were qualitatively classified based on the kind of stimulus required into those requiring light touch, movement of the hair, taps, hard taps, or movements of the joints. We also noted if the neuronal responses were qualitatively perceived to be strong or weak responses. Details of the receptive fields and responses of the neurons were noted, and receptive fields were carefully drawn on photographs of the body surface of each monkey.

At the end of the recording sessions, a few reference locations were marked with electrolytic microlesions by passing 10 µA DC current for 10 seconds (Fig. 1C), or by passing 10 µA current as the electrode was withdrawn at the rate of 25 µm/s to produce a continuous lesion along the electrode track. These lesions allowed us to relate the somatotopic map to subdivisions of the VP nucleus in frontal brain sections. The VP nucleus was defined by its cyto- and chemo-architecture (see below).

**Fig. 1.**
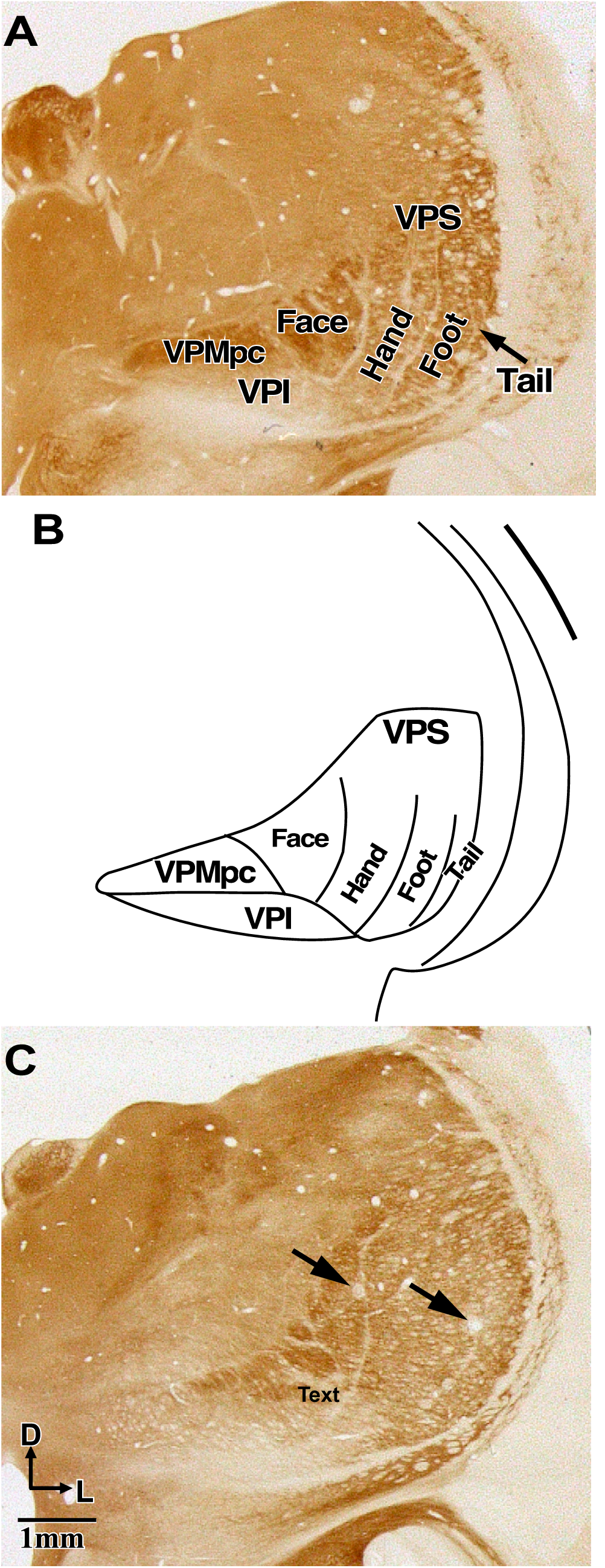
Somatotopy in the ventroposterior nucleus of a macaque monkey. The ventroposterior nuclear (VPN) complex consists of VPM (ventroposterior medial subnucleus), which gets inputs from the face, and VPL (ventroposterior lateral subnucleus), which gets inputs from the hand, foot, trunk and tail in a mediolateral order. VPS (ventroposterior superior nucleus) and VPI (ventroposterior inferior nucleus), which receive deep inputs, are also parts of the VP complex. A. Coronal section of the thalamus through the VPN of a macaque monkey stained for cytochrome oxidase activity. B. An outline of the VPN showing normal somatotopy. VPMpc, parvicellular subdivision of the VP nucleus. C. A coronal section of the thalamus of monkey 98-18MK through the VPN showing electrolytic lesions (arrows) made to align the recording sites in the nucleus (see Methods). The tail subnucleus of VPL is not visible in this section.

During the mapping procedure anaesthesia was continually monitored by monitoring heart rate, responses to toe pinch, and eye blink responses. Supplemental dosage of ketamine and xylazine (20% of the initial dosage, IM or IV) and urethane (250mg/kg) were given as needed to maintain a stable level of anaesthesia. Dexamethasone (2 mg/kg, i.m.) was given to prevent brain swelling. Fluids were continuously given by intravenous route; and body temperature was maintained as described above. The heart rate and blood oxygenation level were monitored throughout the experiment.

### Histology

At the end of the mapping experiments, the animals were deeply anesthetized with pentobarbital and perfused transcardially with 0.1 M phosphate buffered saline (PBS, pH 7.4), followed by 2-3% paraformaldehyde in phosphate buffer (PB), and finally with buffered paraformaldehyde with 10% sucrose. The brain was removed, the thalamus was separated from the surrounding tissue, blocked, and cryoprotected in 30% sucrose. The thalamus was cut in a coronal plane in 40 µm or 50 µm thick sections. Series of sections were stained for Nissl substance (Jain et al., 2008) or cytochrome oxidase activity (Wong-Riley, 1979) to determine borders of the thalamic nuclei.

The spinal cord was removed from the vertebral column. During the dissection level of the lesion was again confirmed by counting dorsal roots from the rostral end. The spinal cord was cryoprotected in the sucrose solution as described above, blocked, and cut in a horizontal plane at a thickness of 50 or 60 µm. The sections were mounted on slides and examined under dark-field to reconstruct the extent of the spinal cord lesion. Blocks from the spinal cord rostral and caudal to the lesion were cut in a coronal plane at a thickness of 50 µm. Series of coronal sections were processed for Nissl substance or cytochrome oxidase activity.

### Reconstruction of the spinal cord lesion site

The extent of the dorsal column lesion was reconstructed from the horizontally cut spinal cord sections and projected onto a transverse view (insets in Fig. 2B, 3B, 4A; also see Jain et al., 2025b; Qi et al., 2010). In order to accomplish this, drawings of the spinal cord sections in the region of the lesion were made using a dark-field view of the microscope (Nikon E800) equipped with a drawing tube, or drawn on the digital photographs of the spinal cord sections acquired using a digital camera attached to the microscope. Maximal extent of the lesion and boundaries of the white matter and the grey matter were drawn for each section, and their dimensions were used to reconstruct the lesion extent in a coronal view.

**Fig. 2A,B.**
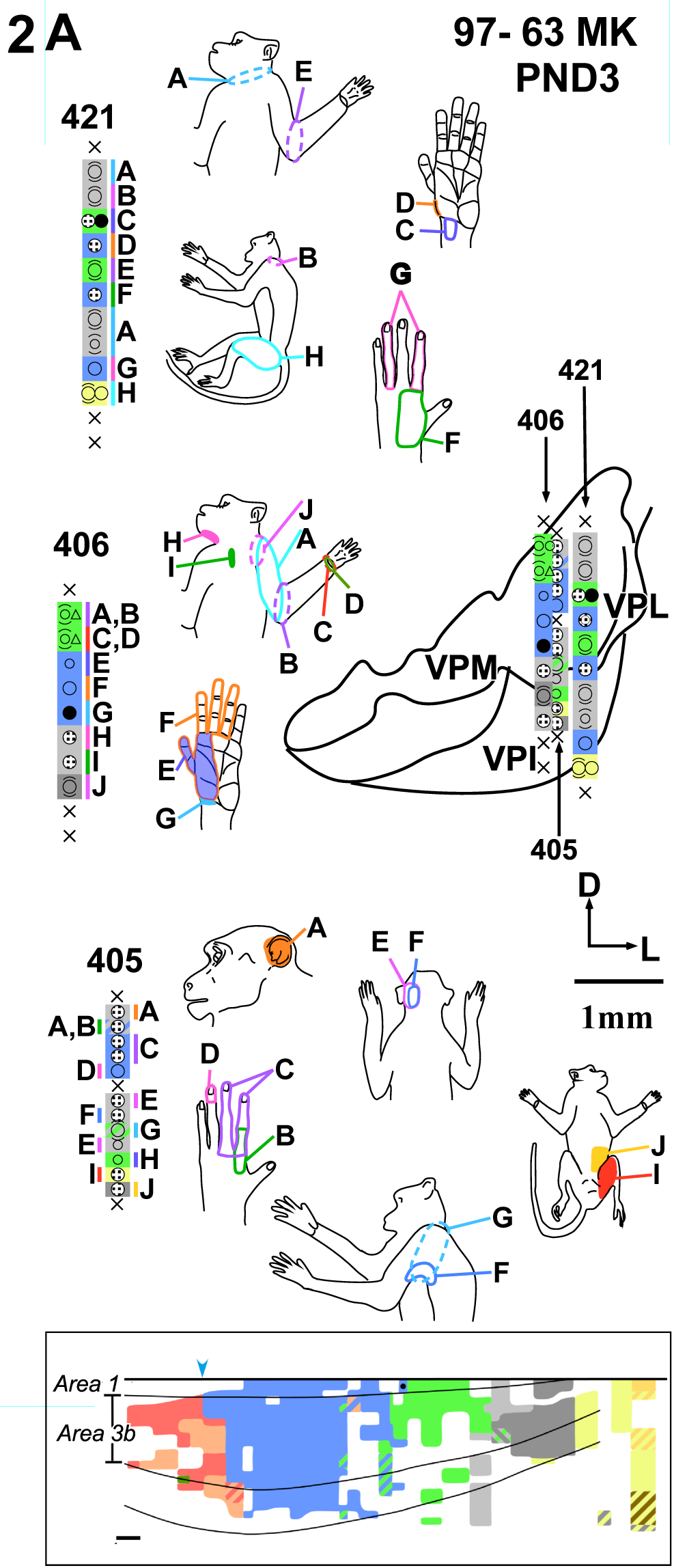
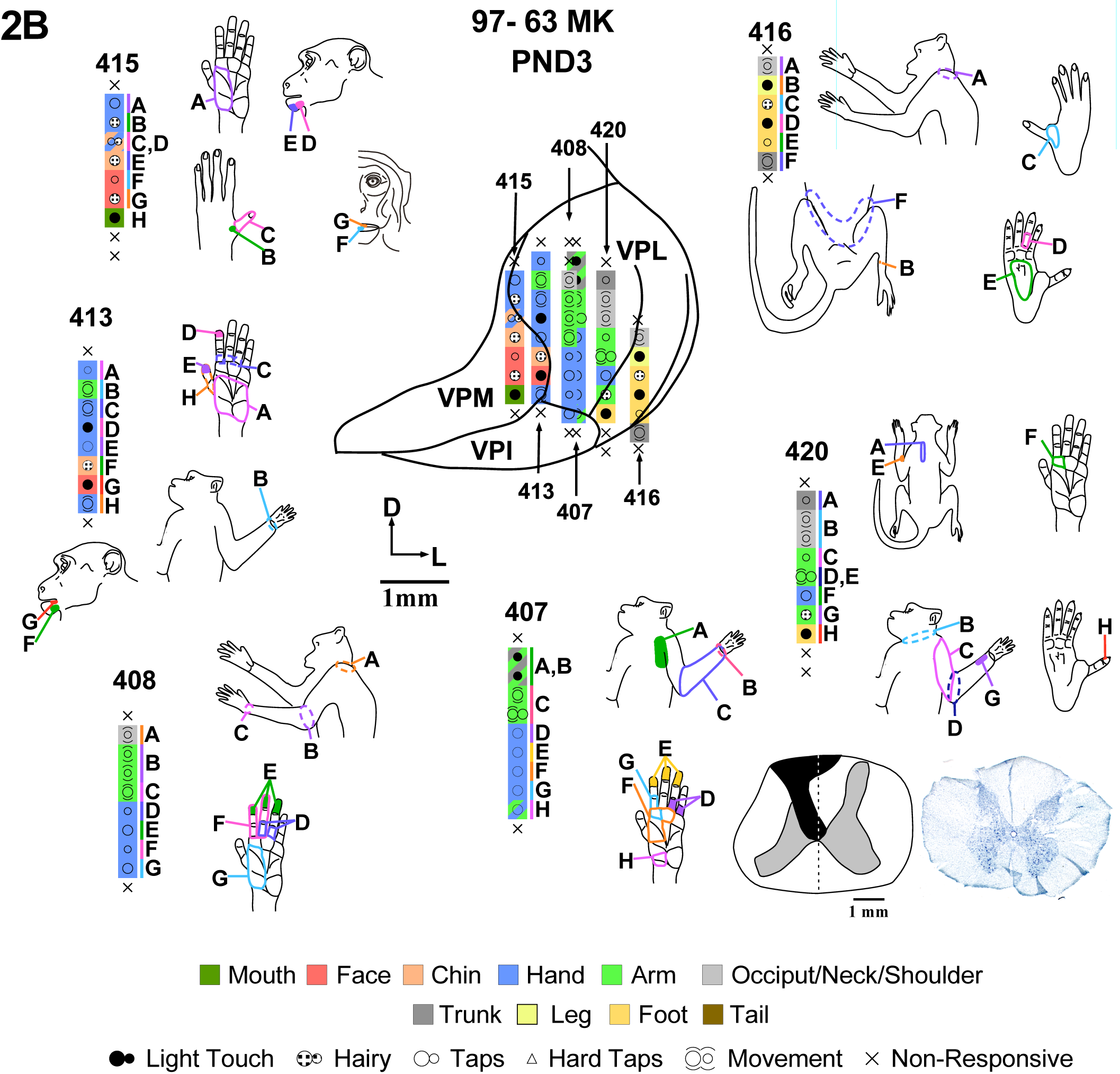
Somatotopy and receptive fields of neurons in the ventroposterior nucleus of the thalamus of monkey 97-63MK that received unilateral dorsal column lesion on PND 3. Inputs from all the body parts are present at the expected somatotopic locations, i.e., the face in VPM, and the hand, foot, and trunk in VPL. The receptive field locations of the neurons encountered in different electrode penetrations through the ventroposterior (VP) nucleus are shown projected on outline drawings (A) a caudal and (B) a rostral coronal section through the VP nucleus. The figurines show receptive field locations and relative sizes at the recording site locations marked with alphabets in different electrode penetrations (numbered). Key to the colors and symbols is shown at the bottom of Fig. 2B. Larger dots indicate a strong response and smaller dots a weak response to the same kind of stimulation as shown in the key. Receptive fields outlined by dashed lined in the figurines are movement receptive fields. Stripes of two colors indicate receptive fields located on two different body parts. Inset in Fig. 2B shows the extent of the spinal cord lesion reconstructed from the horizontal section. The extent of the lesion is colored black; grey is grey matter, and white is white matter. Adjacent photomicrograph shows a Nissl stained coronal section of the spinal cord taken from a level rostral to the lesion. Note shrinkage of the dorsal columns on the left side. Boxed inset at the bottom in Fig. 2A shows for ready reference a simplified version of the somatotopy in area 3b and caudal part of area 1 of this monkey. These data have been previously published in detail (Jain et al., 2025b). The blue arrowhead at the top marks the expected location of the hand-face border. The color key is same as shown in Fig. 2B. For the cortical map medial is to the right and caudal is towards the top; scale bar is 1.0 mm. D, dorsal; L, lateral.

**Fig. 3A-C.**
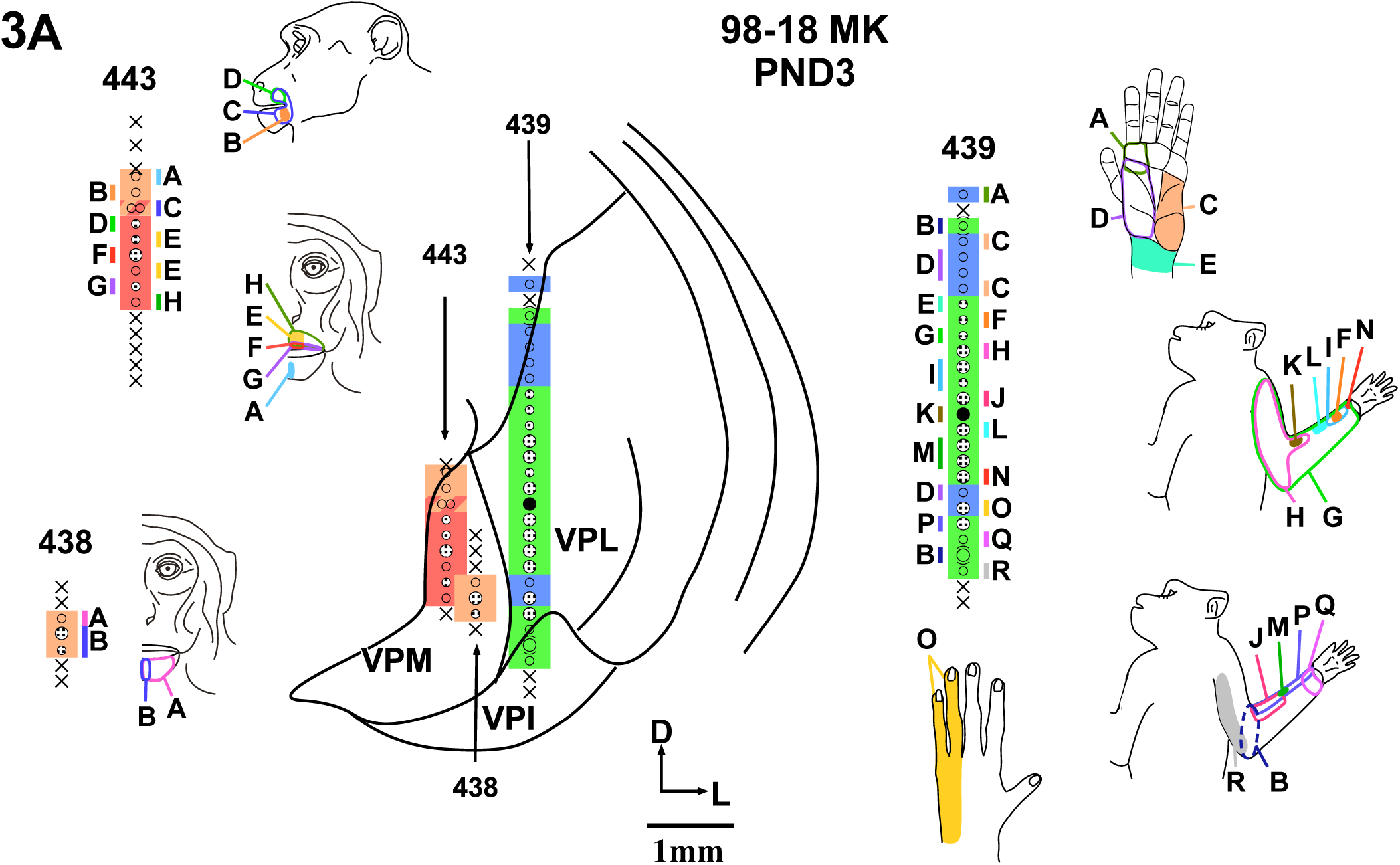
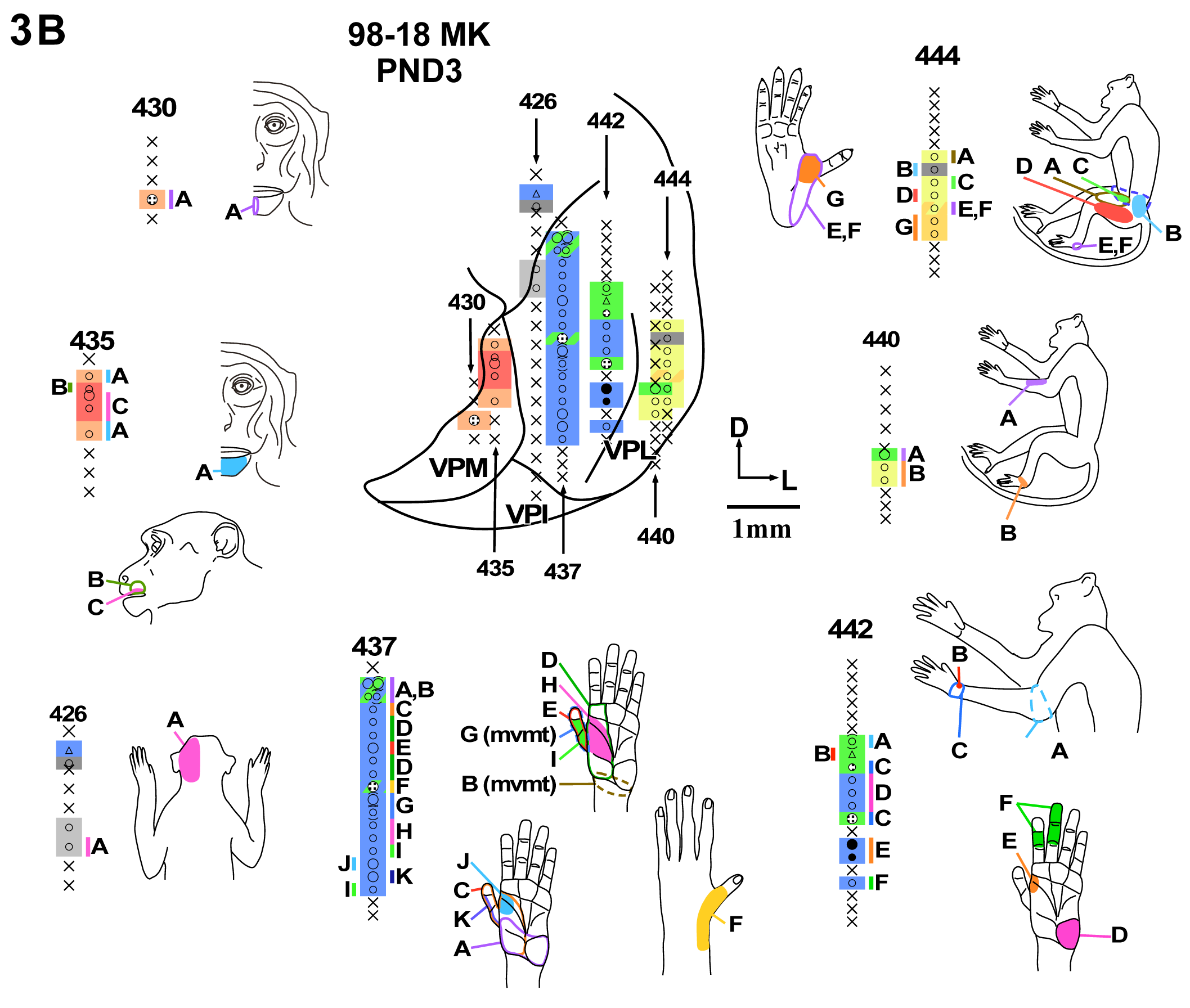
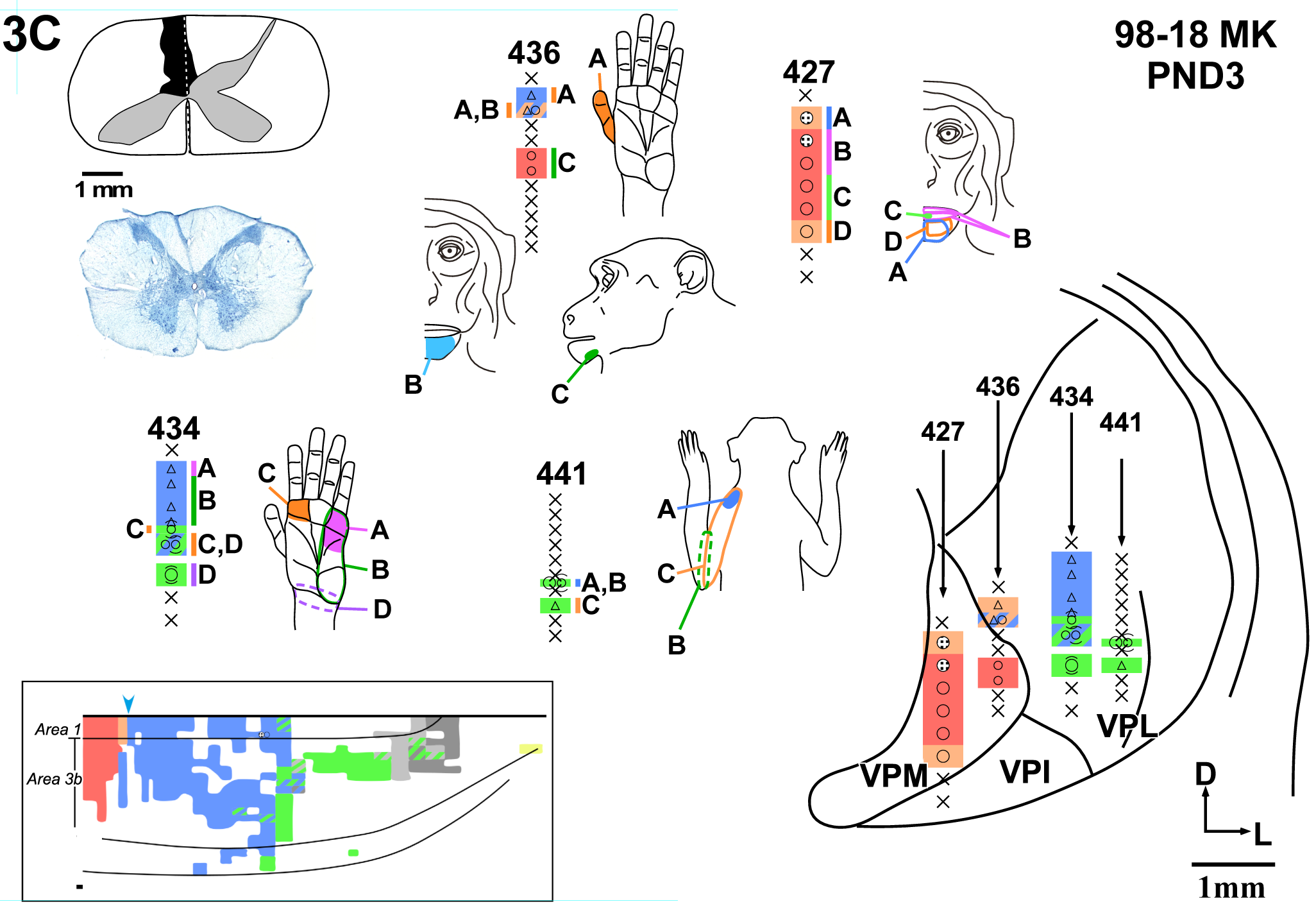
Somatotopy and receptive fields of neurons in the ventroposterior nucleus of the thalamus of monkey 98-18MK that received unilateral dorsal column lesion on PND 3 are shown projected on outline drawings of three (A) caudal to (C) rostral coronal sections through the VP nucleus. All the body parts are represented in the expected somatotopic locations, the face in VPM, and the hand, foot, trunk and tail in VPL. Inset in Fig. 3C shows the extent of the spinal cord lesion and a photomicrograph of a Nissl stained coronal section of the spinal cord taken from a level rostral to the lesion. Boxed inset at the bottom in Fig. 3C shows simplified version of the somatotopy in area 3b and caudal part of area 1 of this monkey (see Jain et al., 2025b for details). Other conventions as in Fig. 2.

**Fig. 4A-D.**
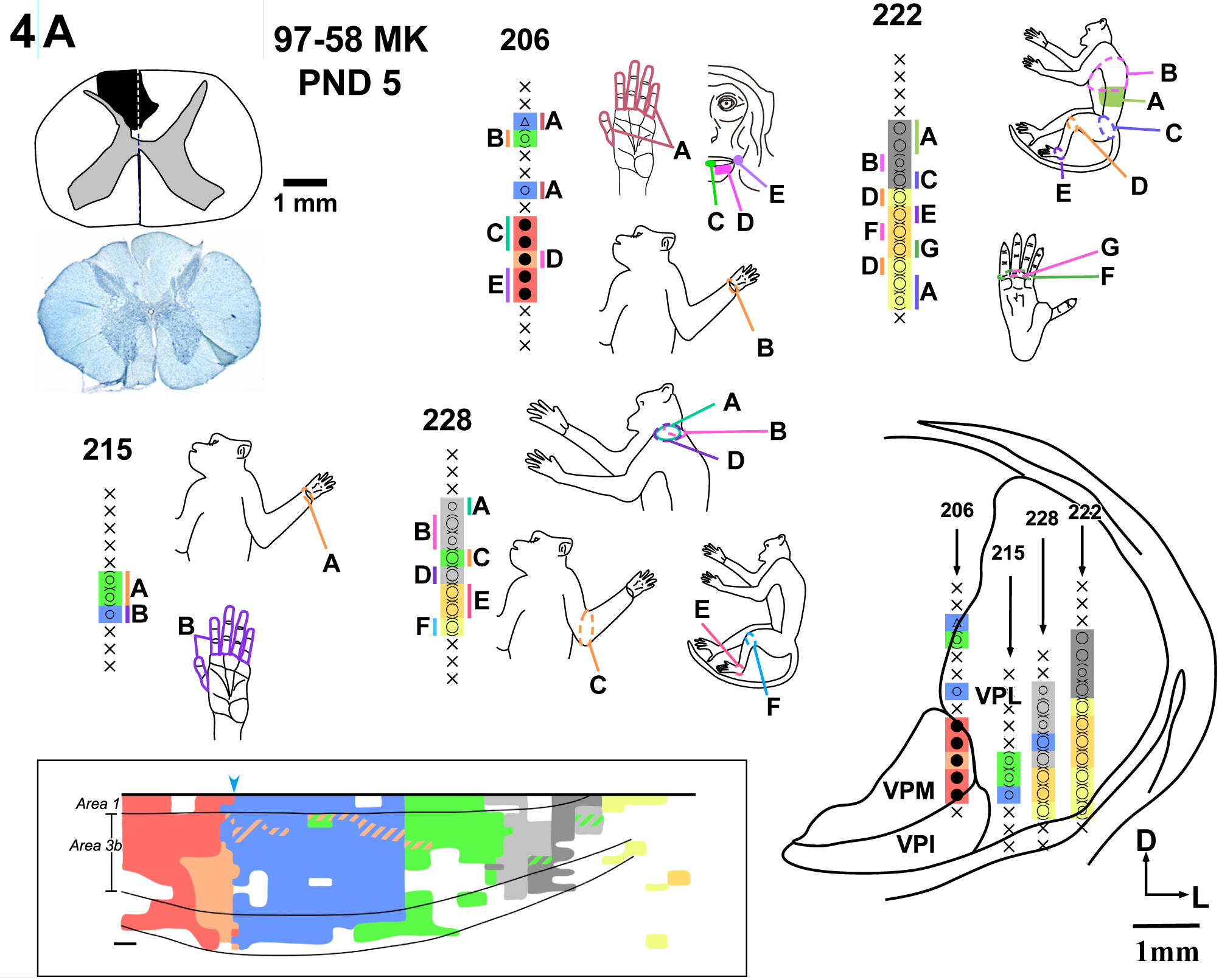
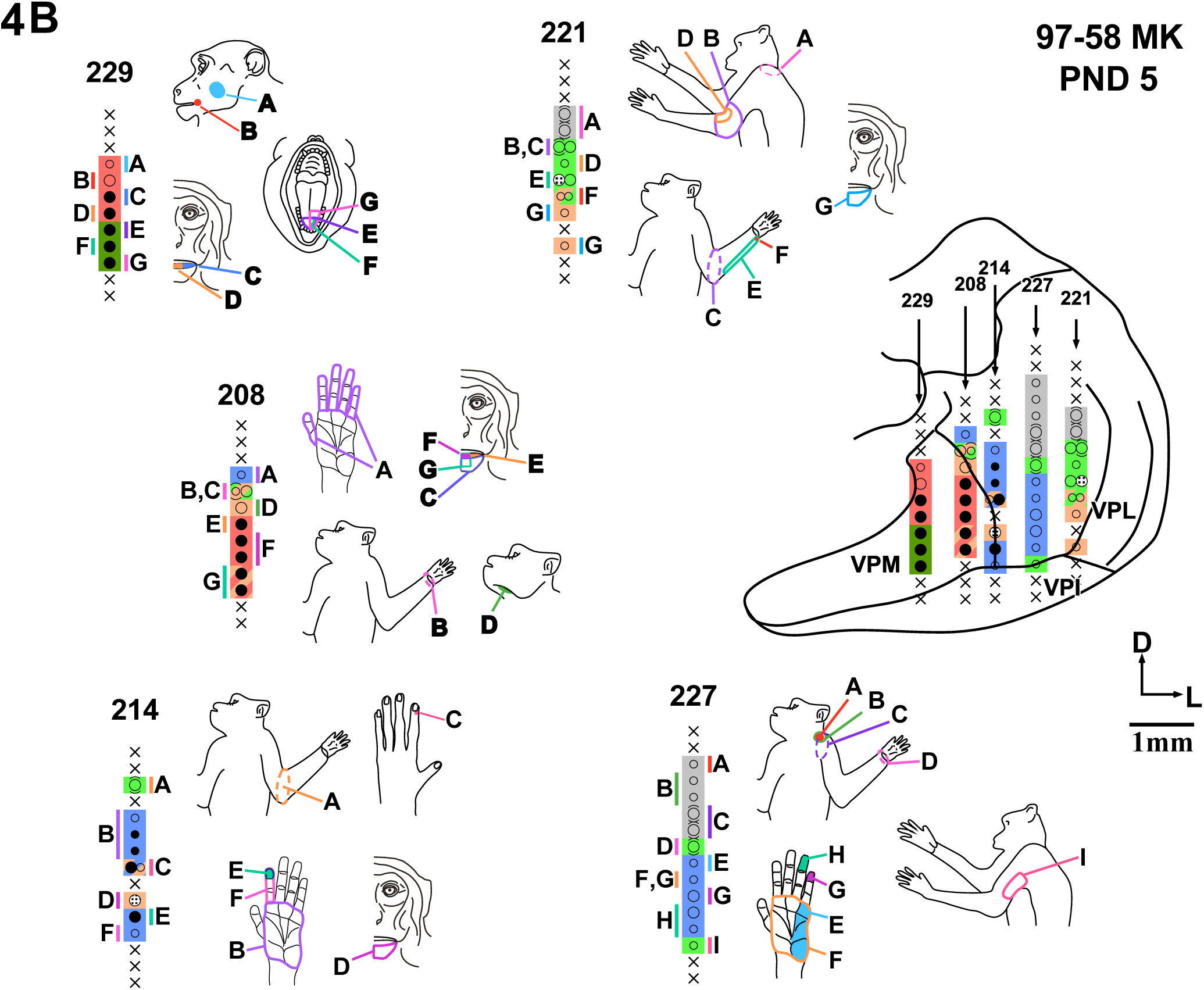
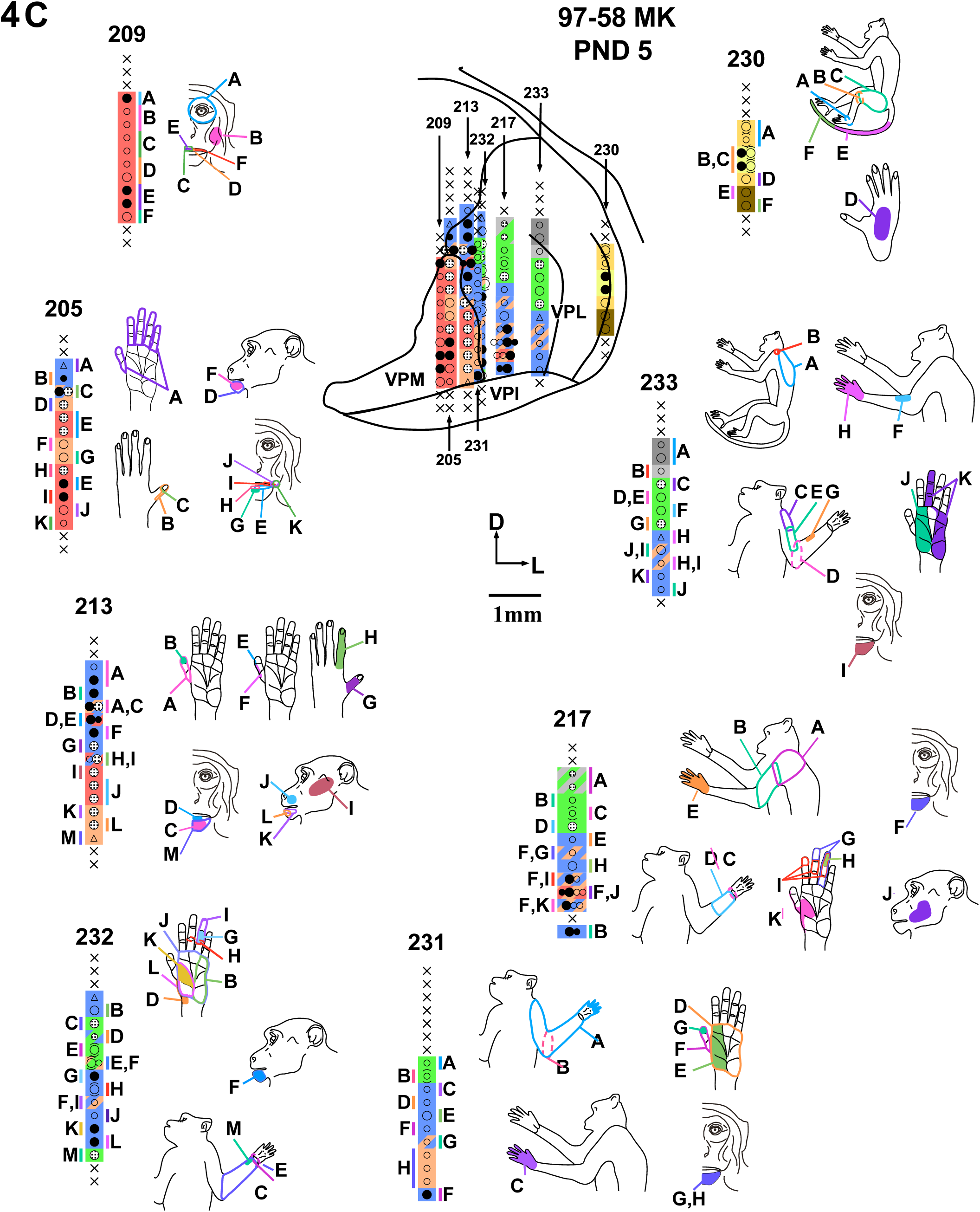
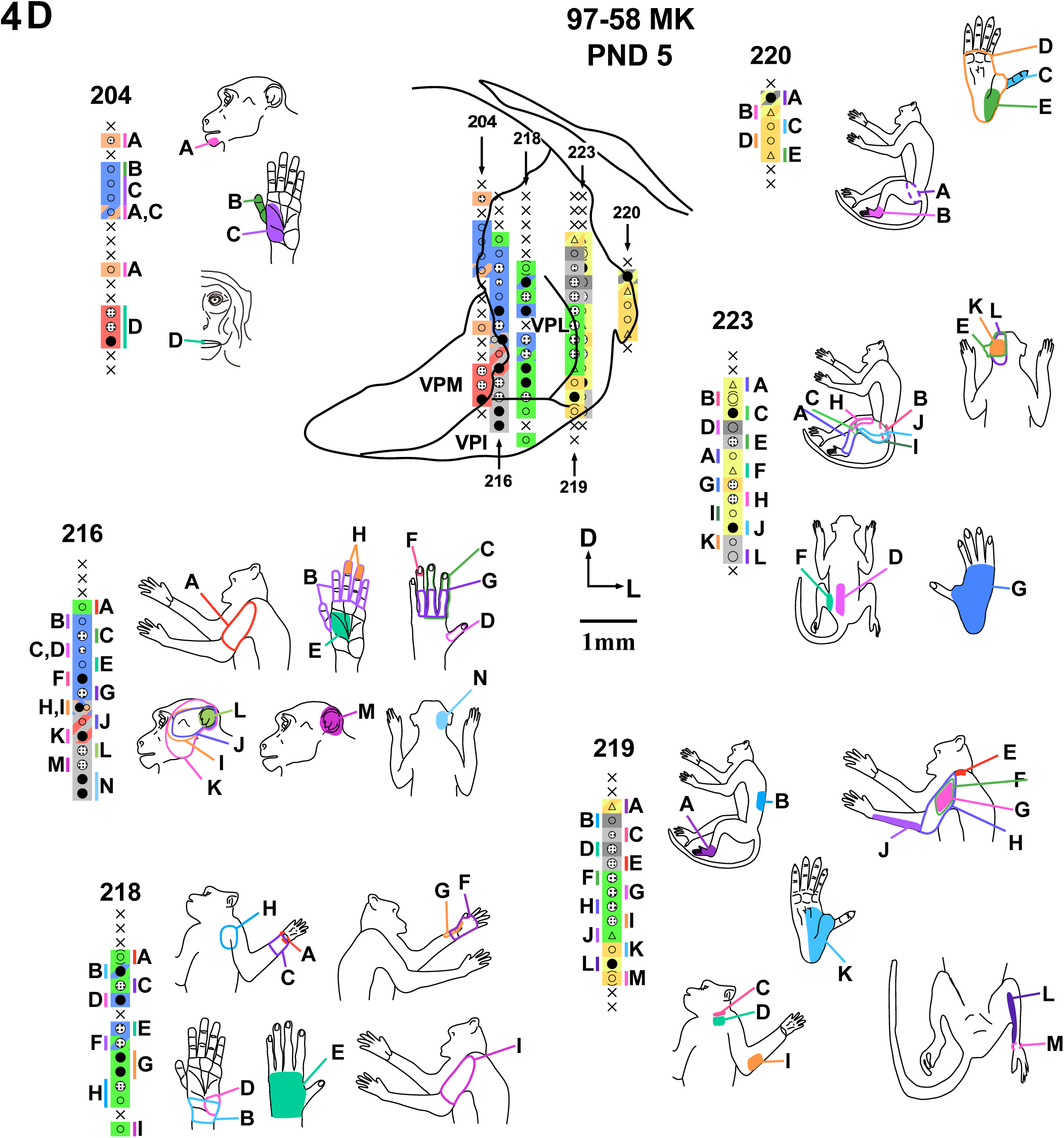
Somatotopy and receptive fields of neurons in the ventroposterior nucleus of the thalamus of monkey 97-58MK that received unilateral dorsal column lesion on PND 5 are shown projected on outline drawings of four (A) caudal to (D) rostral coronal sections through the VP nucleus. All the body parts are represented in the expected somatotopic locations, the face in VPM; and the hand, foot, trunk, and tail in VPL. Note that at a few recording sites in VPL, the receptive fields are on the chin. Inset on top left in Fig. 4A shows extent of the spinal cord lesion and a photomicrograph of a Nissl stained coronal section of the spinal cord taken from a level rostral to the lesion. Boxed inset at the bottom left in Fig. 4A shows simplified version of the somatotopy in area 3b and caudal part of area 1 of this monkey (see Jain et al., 2025b for details). Other conventions as in Fig. 2.

### Reconstruction of the somatotopic maps of the VP nucleus

Somatotopic maps were reconstructed using Canvas software (ACDSee, WA, USA) as described in detail elsewhere (Jain et al., 2025b; Tandon et al., 2009). Neuronal responses requiring light touch, hair stimulation, taps, or hard taps were represented by different symbols and marked accordingly. Response to each kind of stimulus, whether it was strong or weak was also indicated by the size of the symbol. Receptive field locations were coded by different colours. All the reconstructed electrode tracks were placed in their relative positions and mapped onto the nearest section of the thalamus through the VP nucleus. Boundaries of the VPM (ventroposterior medial) and VPL (ventroposterior lateral) subnuclei, and the adjacent nuclei were identified in sections stained for Nissl substance or CO activity and drawn (Fig. 1). Different body part representations were coded by different colours to illustrate location of the receptive fields for each penetration and drawn on the corresponding section to visualize the somatotopy in the entire rostro-caudal and medio-lateral extent of the VP nucleus. The receptive fields were drawn on the figurines of the monkey body parts for ease of visualization. Although a large extent of the thalamus was explored as described above, in order to avoid clutter we have not illustrated penetrations where neurons were non-responsive to any peripheral stimulation because the track did not pass through the VP nucleus. In addition, to indicate the boundaries of the responsive region we have shown only one or few non-responsive recording sites dorsal and ventral to the recording sites where neurons responded to peripheral stimulation. However, tracks with non-responsive sites that passed through the VP nucleus, and non-responsive sites that were intervening between responsive sites are illustrated.

## RESULTS

We made unilateral lesions of the dorsal columns of the spinal cord at cervical levels in macaque monkeys when they were between 3 and 9 days old. The ventroposterior nucleus (VPN) of the thalamus was mapped when the monkeys were between 3-5 years old (Table 1). Results from each monkey are described in the order of increasing age at the time of the lesion.

### Monkey 97-63MK, lesioned on PND 3

Three days after birth, the left dorsal columns of monkey 97-63 were transected at C5 level (Table 1). Reconstruction of the lesion site revealed that although most of the dorsal column fibres were severed (Fig. 2B, inset), there was small sparing ventromedially. The fasciculus cuneatus appeared to be nearly completely transected.

When the monkey was 3 years and 4 months old (Table 1), somatotopy of the contralateral VPN was determined (Fig. 2A, 2B). We made a total of 13 penetrations through the VPN. The overall somatotopy was similar to what has been described before for normal monkeys (Jones and Friedman, 1982; Kaas et al., 1984; Mountcastle and Henneman, 1952). The hand representation was medial-most in VPL (ventroposterior lateral subnucleus). The arm representation was located lateral and dorsal to the hand representation. Dorsal VPL had neurons responding to touch on the occiput, neck, shoulder, and trunk. Lateral to the forelimb representation, inputs from the hindlimb were encountered as in normal animals (Fig. 2B, Pen #416 and 420). Two of the penetrations (Fig. 2B, Pen #413 and 415) passed through the VPM (ventroposterior medial subnucleus), where we encountered neurons responding to the touch on the chin and face.

Although the somatotopy was normal-like in the VPL, the receptive fields were generally larger than expected (e.g. Fig. 2A, Pen# 406 sites E and F; Fig. 2B, Pen# 413 site A, #408 site G, Pen # 407 all RF on the glabrous hand). Throughout the text normal-like implies comparison to data from normal monkeys mapped in our laboratory under similar conditions (see Tandon et al., 2009; Halder et al., 2018; Jain et al., 2008). For example, the receptive fields extended over multiple digits (Pen# 408, sites D, E, F; #406, site F) or the entire palm (Pen #413, site A). Moreover, at a large number of the recording sites, neurons responded weakly to the touch, required light taps, or movements of the joints for evoking responses (refer to the symbol key given in Fig. 2B). These receptive field properties resembled those in the VPI (ventroposterior inferior nucleus) (Pen #406, 421).

In the lateral-most penetrations through the hindlimb subnucleus of the VPL, the neurons responded vigorously to the touch on the toes or the plantar foot (Fig. 2B, Pen# 416 site B, C, D, and #420 site H). These responses were likely due to sparing of the fibres in the fasciculus gracilis (see inset in Fig. 2B). At the recording sites located in VPM the neurons responded vigorously to touch on the lips or hairs on the chin (Fig. 2B, Pen# 413 and 415). These receptive fields were as small as those in the normal animals. Within VPL and VPM, we did not find any region where neurons were unresponsive to peripheral stimulation.

We have previously shown that in this monkey the somatotopy in area 3b is normal like (Fig. 1 of Jain et al., 2025b, also see inset in Fig. 2A). Thus, if lesions of the dorsal columns are made 3 days after birth, somatotopy in VPN and the somatosensory cortex are similar and very normal-like. In VPL, particularly in the hand subnucleus, neurons had large receptive fields and responded weakly to taps or movements of the body parts rather than to a light touch.

### Monkey 98-18MK, lesioned on PND 3

Three days after birth, the left dorsal columns of this monkey were transected at C3/C4 level (Table 1). Reconstruction of the lesion site revealed that the lesion was complete (Fig. 3C, inset). Both the fasciculus cuneatus and fasciculus gracilis were completely transected. The dorsal horn as well as the grey matter up to the level of the central canal were not discernible.

When monkey 98-18MK was four years old (Table 1), somatotopy of the contralateral VPN was determined (Fig. 3A-C). In this monkey, 17 penetrations were made through VPN. The overall somatotopy in VPN was normal-like as it was for the monkey 97-63MK described above. Neurons responding to the trigeminal nerve inputs were present in VPM (Pen # 427, 430, 435, 436, 438, 443), and they had small receptive fields. More laterally, neurons responded to the tactile stimulation of the hand and arm in the medial part of VPL. At most of these recording sites, the neurons responded weakly and required taps rather than light touch to evoke responses (Fig. 3B and 3C). The receptive fields were large, often covering an entire digit, multiple digits, or a pad of the palm (Fig. 3A, Pen# 439; Fig. 3B, Pen#437, 442; Fig. 3C, Pen# 434, 436). Interestingly, for the receptive fields located on the anterior forearm, the receptive fields were small, and the neurons responded vigorously to the light touch on hairs (Fig. 3A, Pen #439). The activating forearm inputs likely enter the spinal cord rostral to the site of the lesion (Florence et al., 1988).

Unlike neuronal response in monkey 97-63MK described above, neurons in this monkey responded weakly to taps on the lower limb (Fig. 3C, Pen #440 and 444), reflecting a complete lesion of the contralateral dorsal columns. Finally, at many recording sites, neurons did not respond to touch on any part of the body (e.g., Pen# 426, #442).

Thus, in monkey 98-18MK with the lesion of the dorsal columns at 3 days after birth, the somatotopy in VPN and area 3b of the cortex (see Fig. S2 of Jain et al., 2025b, also see inset in Fig. 3C) was similar and normal-like. However, unlike normal monkeys, neurons in the VPL had large receptive fields, and they responded weakly and required taps on the skin rather than light touch to evoke responses.

### Monkey 97-58MK, lesioned on PND5

Left dorsal columns of monkey 97-58 were transected at C3/C4 spinal level, 5 days after birth (Table 1). The lesion was nearly complete. Reconstruction of the lesion site showed sparing of fasciculus cuneatus in the dorsolateral part near the dorsal horn, and in the ventral-most region (Fig. 4A, inset). Contralateral VPN was mapped when the animal was just over 4 years of age (Fig. 4A-D).

Neurons at recording sites in VPL responded to tactile stimulation of the hand, arm, occiput, neck, shoulder and trunk as expected (Fig. 4A-D). At most of the recording sites in VPL neurons did not respond to the light cutaneous touch, but required taps or movements of the body parts to evoke a response (Fig. 4 A-D, see key to the symbols in Fig. 2B). The receptive fields were often large. The receptive fields covered an entire digit (e.g., Fig. 4B, Pen#214, site F), multiple digits (e.g., Fig. 4C, Pen#205, site A), or the entire palm (e.g., Fig. 4B, Pen#214, site B; Fig. 4C, Pen#232, site J). In addition, neurons at several recording sites in the hand subnucleus of VPL responded to touch on the chin (Fig. 4B, Pen #221; Fig. 4C, Pen # 217, 233), which is unlike for the two monkeys lesioned on PND3 described above. At some of the recording sites, neurons had receptive fields on both the hand and the arm (e.g., Fig. 4C, Pen# 233, sites H, I, and J). More laterally, neurons responded to touch on the lower limb or tail in the hindlimb and tail sub-nuclei of VPL (Fig. 4C, Pen #230; Fig. 4D, Pen # 220, 223). Finally, for a few recording sites in VPL, neurons did not respond to touch on any part of the body.

Neurons at recording sites in the VPM had small receptive fields and responded vigorously to touch on the skin of the lips, or hairs on the chin or other parts of the face (e.g., Fig. 4a, Pen #206; Fig. 4B, Pen#208, 229; Fig. 4C, Pen# 205, 209, 213; and Fig. 4D, Pen# 204). In this monkey, we also encountered recording sites in the expected location of the medial part of VPM where neurons responded to tactile stimulation of the oral cavity structures (Fig. 4B, Pen# 229).

Thus, in VPL of this monkey, most of the somatotopy was normal-like, at few recording sites where receptive field on the hand were expected, neurons responded to touch on the chin. The somatotopy was similar in both VPN and the somatosensory cortex (see Fig. 3 of Jain et al., 2025b, also see inset in Fig. 4A).

### Monkey 99-84MK, lesioned on PND 9

Dorsal columns of this monkey were lesioned on the left side at C3/C4 spinal level on the 9^th^ day after birth. The contralateral VPN was mapped when the animal was about 5 years old (Table 1). The lesion appeared to be almost complete. Reconstruction of the lesion site from spinal cord sections was not possible due to histological difficulties. However, coronal sections of the spinal cord rostral to the lesion revealed that the dorsal columns were asymmetrical with a considerable reduction in size ipsilateral to the lesion, suggesting a nearly complete lesion (Fig. 5 inset; this figure is the same as shown in (Qi et al., 2010).

**Fig. 5A,B.**
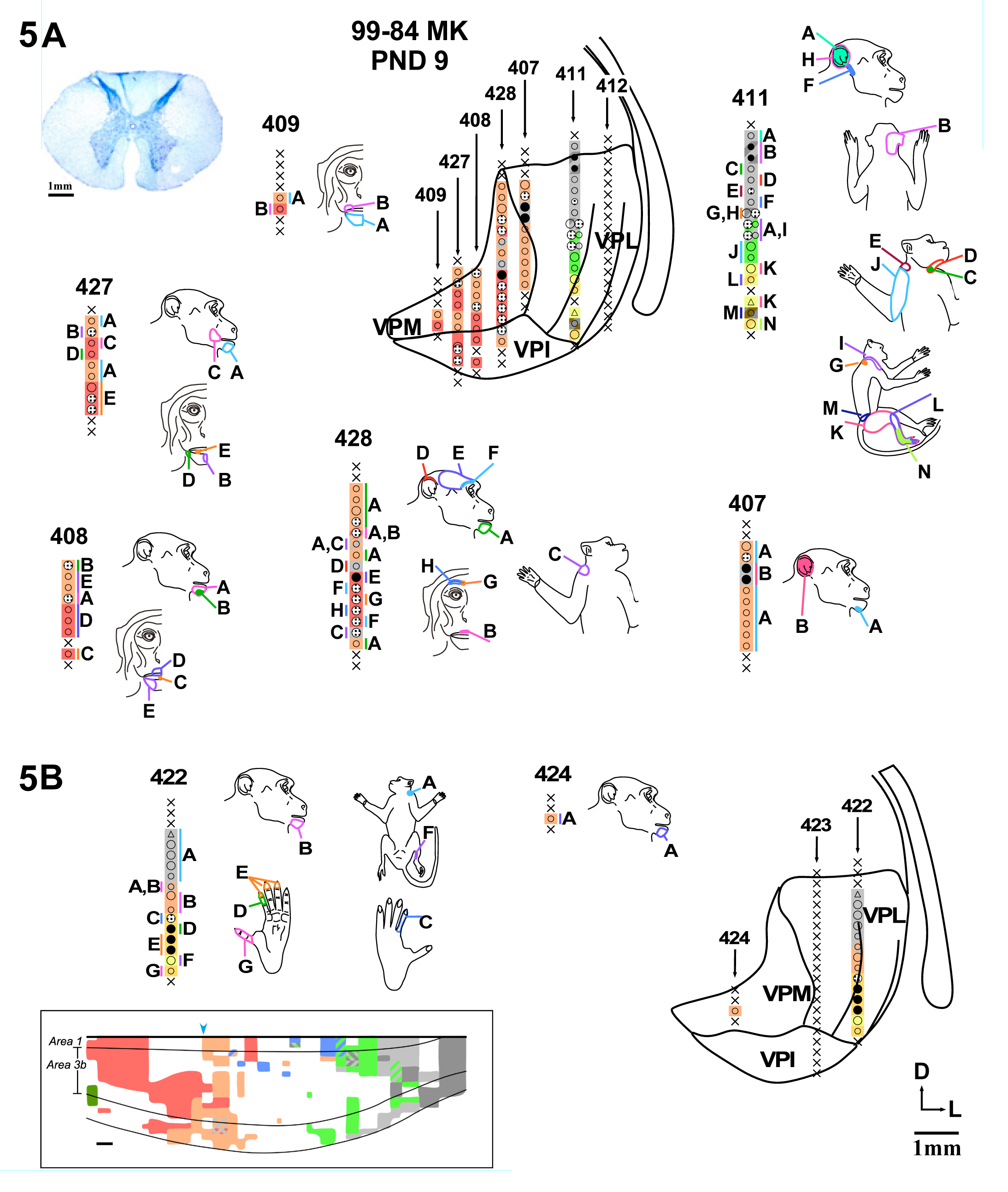
Somatotopy and receptive fields of neurons in the ventroposterior nucleus of the thalamus of monkey 99-84MK that received unilateral dorsal column lesion on PND 9 are shown projected on outline drawings of (A) a caudal and (B) a rostral coronal section through the VP nucleus. The somatotopy is normal in VPM. In VPL at a few locations, the receptive fields are on the chin, whereas at many locations, neurons do not respond to touch on any part of the body. The inset shows a section of the spinal cord taken rostral to the site of the lesion and stained for Nissl substance. Note the considerable shrinkage of the dorsal columns on the lesioned left side, suggesting a complete or nearly complete lesion of the dorsal columns. Boxed inset at the bottom in Fig. 5B shows simplified version of the somatotopy in area 3b and caudal part of area 1 of this monkey (see Jain et al., 2025b). Other conventions as in Fig. 2.

We encountered neuronal responses in 10 of the 14 penetrations made through the VPN (Fig. 5). In VPL, we did not find any neurons that responded to touch on the hand. Neurons with receptive fields on the arm were encountered at four recording sites in one penetration (Fig. 5A, Pen# 411). In this penetration neurons at other recording sites responded to touch on the occiput and neck dorsally, and the lower limb and base of the tail ventrally. At a few recording sites, neurons responded to touch on the chin. For example, in the penetration #422 (Fig. 5B), which was through the hand subnucleus of VPL, the neurons responded to tactile stimulation on the shoulder dorsally and chin ventrally. In the same penetration, neurons responded to tactile stimulation of the foot in the ventral most part as the electrode entered the lower limb subnucleus of VPL. At several recording sites through the VPL, neurons did not respond to touch on any part of the body (Pen #412, #423). We also made many penetrations through the VPM in this monkey (Pen# 407, 408, 409, 427, 428). At all these recording sites in VPM, neurons responded to touch on the face as expected.

In the expected hand region of area 3b of this monkey, the neurons at most of the recording sites did not respond to touch on any part of the body (see Fig. 4 of (Jain et al., 2025b). In the lateral part of the expected hand representation om area 3b, there was a medial-ward expansion of the chin representation. Thus, the somatotopy in VPN and the somatosensory cortex (see Fig. 4 of Jain et al., 2025b, also see inset in Fig. 5B) of this monkey was similar, with large regions where neurons did not respond to touch on any part of the body, and a few recording sites where neurons responded to tactile stimulation of the chin.

Thus, in all the four monkeys with unilateral lesions of the dorsal columns at cervical levels made when the animals were between 3-9 days old, the somatotopy in area 3b reflected that in the VPN. The somatotopy was normal-like in monkeys with lesions made on PND3 and PND5. However, the receptive fields on the hand were larger than expected, and the neuronal responses were weak, often requiring taps on the skin to evoke responses rather than a light touch. In the monkey with a lesion made on PND9 the neurons were unresponsive to touch on any part of the body in large expanse of VPL where inputs from the hand are normally expected. At a few sites in the deafferented VPL neurons responded to touch on the chin.

## DISCUSSION

We determined somatotopic organization of the ventroposterior nucleus (VPN) of the thalamus in four monkeys that sustained unilateral lesions of the dorsal columns of the spinal cord at cervical levels soon after birth. The lesions were made on postnatal day 3 (PND3) in two monkeys, on PND5 in one monkey, and on PND9 in the fourth monkey. The monkeys were mapped when they were between 3 and 5 years of age. Our major finding is that in monkeys with lesions done on PND3 or PND5, somatotopy in the ventroposterior lateral nucleus (VPL) was normal-like, whereas in the monkey that was lesioned on PND9, most of the deafferented neurons in the VPL did not respond to touch on any part of the body. In this monkey neurons at a few recording sites respond to touch on the face. In the visual system, inactivation of the visual inputs early in development results in an irreversible loss or degradation of vision from the deprived eye inputs even if these inputs are restored after what is called the ‘critical period’ has passed (Blakemore and Van Sluyters, 1974; Hubel et al., 1977; LeVay et al., 1980). In the somatosensory system, there appears to be a reverse critical period because early loss of tactile inputs via the dorsal columns up to PND5 results in normal-like topography in the somatosensory cortex, while a lesion done after PND9 results in irreversible and abnormal reorganization (Jain et al., 2025b, and this study).

### Deafferentation induced reorganization is similar in the VPN and the somatosensory cortex

We have previously reported on reorganization of the somatosensory cortex in the same monkeys (Jain et al., 2025b). In the monkeys with lesions done on PND3 and PND5, the cortical areas 3b, 1, and 2 had normal-like somatotopic reorganization, as described here for VPN. In the somatosensory cortex and VPL of the monkey lesioned on PND5, there were a few recording sites where neurons responded to touch on the chin. In the monkey lesioned on PND9, in both the somatosensory cortex and VPN, neurons in the deprived regions either did not respond to touch on any part of the body (see Results) or at a few sites responded to touch on the face. Thus, in each of the four monkeys the somatotopy in the somatosensory cortex and VPN was similar. We have previously reported that in adult monkeys with deafferentation due to lesions of the dorsal columns, reorganization of the VPN and the cortical area 3b are similar (Halder et al., 2018; Jain et al., 2008). Post-lesion reorganization has also been found to be similar in the VPN and the somatosensory cortex after other kinds of peripheral deafferentations (Garraghty and Kaas, 1991; Kaas et al., 1999; Merzenich et al., 1983). This likely reflects high fidelity of the somatotopy of the projections from VPN to the somatosensory cortex (Nelson and Kaas, 1981), so that any reorganization in the VPN is reflected in the somatosensory cortex. Yet, cortical mechanisms also appear to play a role in shaping the post-lesion reorganization of the cortex (Florence et al., 1996; Halder et al., 2018). Following spinal cord injuries in adult animals neurons in the deafferented parts of the VPL respond to touch on both the chin and inputs from the occiput/neck/shoulder. Similar reorganization was observed in the cuneate nucleus of the brainstem (Halder et al., 2018; Kambi et al., 2014). However, following lesions in adult animals, in area 3b only the chin inputs reactivate neurons in the deafferented hand cortex; inputs from the occiput/neck/shoulder activate neurons only in the medial-most regions of the deprived hand cortex (Jain et al., 2008). We proposed that there is a resorting of ascending inputs from the VPN to area 3b such that reorganization in the VPN is not passively reflected in the somatosensory cortex.. Here also in the monkey 99-84MK that was lesioned on PND9, there appears to be more extensive expansion of the occiput and shoulder representation in the VPL than was apparent in the cortex (see, Jain et al., 2025b).

### Sources of somatotopic inputs in monkeys with lesions done at PND3 and PND5 and mechanisms of reorganization

Since dorsal column lesions appeared to be complete or nearly complete in the monkeys lesioned on PND3 or PND5, the only possible source of somatically appropriate activation of VPL is by inputs via alternate spinal pathways such as the lateral and ventral spinothalamic tracts (STT). The lateral SST largely conveys nociceptive inputs, whereas the lamina V neurons of ventral STT have a wide dynamic range that respond to light touch, pressure, noxious mechanical and thermal stimuli, as well as joint and muscle stimulation (Apkarian and Hodge, 1989; Willis et al., 1979). We found that neurons in the VPL of monkeys with dorsal column lesions made on PND3 and PND5 had relatively large receptive fields, sometimes extending to multiple digits, and required taps or movements of the body parts to evoke responses. In area 3b of these monkeys at most of the recording sites in the deprived cortex, the neurons responded to taps on the skin (Jain et al., 2025b). In normal monkeys, the receptive fields of the neurons in the VPL and area 3b are small, and neurons respond vigorously to light touch on the skin or hair (Felleman et al., 1983; Kaas et al., 1984).

In adult monkeys, both dorsal column-medial lemniscus (DC-ML) pathways and STT terminate in VPL, although in separate compartments. Inputs from STT terminate in patches or islands surrounded by DC-ML inputs (Boivie, 1979; Craig, 2008; Shi and Apkarian, 1995). During the early developmental period, there is an extensive overlap of different modality specific inputs at all levels of the somatosensory pathway starting from the dorsal horns (Fitzgerald and Jennings, 1999; Galea and Darian-Smith, 1995). We have proposed that during early development exuberant tactile inputs from the DC-ML pathways and the STT co-terminate in the same territory in VPL (Jain et al., 2025b). During early development, the nature of inputs specifies modality specific representational identities (Pouchelon et al., 2014; Simi and Studer, 2018) and refines the representations (Granmo et al., 2008), leading to the compartmentalisation of pathway specific terminations in adult animals. We suggest that if DC-ML tract is transected by PND5, the exuberant STT pathways do not withdraw to their mature compartments due to a lack of competition from the DC-ML inputs. This retains normal somatotopy even in the absence of the DC-ML afferents although with different receptive field properties. A similar mechanism has been proposed for plasticity of the ocular dominance columns in the primary visual cortex after monocular deprivation early in development (Takahata et al., 2018).

### Comparison with reorganization in adult monkeys

We propose that by PND9, DC-ML and STT pathways attain mature-like compartmentalization of their terminations in VPL, since neurons at a large number of recording sites in both the somatosensory cortex and VPL do not respond to tactile stimulation of any part of the body in monkeys lesioned on PND9 (Jain et al., 2025b), and this study). Therefore, transection of the dorsal columns on the 9^th^ postnatal day or later results in permanent loss of somatotopically appropriate inputs from parts of the body below the site of the lesion. Withdrawal of STT and DC-ML terminations in VPL to their separate compartments likely completes between PND5 and PND9.

Following lesions of the dorsal columns in adult monkeys, there is a large-scale expansion of the representation of face, specifically chin inputs, in the deafferented parts of area 3b of the cortex (Jain et al., 1997; Jain et al., 2008; Tandon et al., 2009) and in the VPL (Halder et al., 2018; Jain et al., 2008). It is not clear why in the animal lesioned at PND9, both VPN and the somatosensory cortex show only limited expansion of the face representation. Lack of a large-scale expansion of the chin representation is also observed in the somatosensory cortex of the monkeys receiving lesions of the dorsal columns on PND11 or PND12 (Jain et al., 2025b). Expansion of the chin representation in monkeys with lesions of the spinal cord sustained as adults is due to somatotopic reorganization in the cuneate nucleus of the brainstem.(Halder et al., 2018; Kambi et al., 2014). The reorganization in the cuneate nucleus is due to the sprouting of the arbors from the adjacent trigeminal nucleus into the deafferented cuneate nucleus (Jain et al., 2000; Kambi et al., 2014). Reorganization of the cuneate nucleus is reflected upstream due to the pre-existing projections from the cuneate nucleus to the VPL, and from VPL to the somatosensory cortex (Halder et al., 2018; Kambi et al., 2014). We do not know if there is any reorganisation in the somatotopy of the cuneate nucleus of the monkeys lesioned on PND9 or later. However, sprouting from trigeminal into the cuneate nucleus is also seen in the animal lesioned on PND9 (Jain et al., 2025b).

Brainstem medullary nuclei in macaque monkeys show an adult-like differentiation as early as E75 (Jain et al., 1995). Afferents from the brainstem nuclei reach the thalamus, and the thalamic afferents reach the somatosensory cortex in prenatal stages (Davidson et al., 2010). Although the thalamocortical connections are anatomically not yet adult-like during early postnatal days (Darian-Smith et al., 1990a, 1990b). In response to a nerve injury sustained at E75 or E114, the deprived regions of the cuneate nucleus show an increase in GAP43, a growth associated protein. This suggests that these deprived regions stay in an active growth modeatleast until 10 weeks after birth, which likely facilitates arbor remodelling and plasticity (Jain et al., 1995). However, if the nerve injury is sustained at PND11, no such increase in GAP43, i.e., there is no reversion to the growth mode in the cuneate nucleus (Jain et al., 1995). We do not know why an extensive expansion of the chin inputs is not seen in either area 3b or the VP nucleus of animals that receive lesion of the dorsal columns between PND9 and PND13. It is possible that the reorganization is more restricted due to a more severe degeneration of the neurons following large-scale deafferentations in early postnatal days (Kutcher and Duffy, 2007; Sesma et al., 1984; Sloper et al., 1988; Takahata et al., 2018).

## Role of Authors

Drs. N.J. and J.H.K. designed research; N.J., H.-X.Q., D.C.L, performed research; N.J. and A.R. analyzed data; N.J. wrote the manuscript; N.J., H.-X.Q., A.R., D.C.L., and J.H.K. edited the manuscript.

## Acknowledgments

This study was supported by NIH grant NS16446 and NS129982 to JHK; NJ was a Wellcome Trust Senior Research Fellow during part of this study (063259/Z/00/Z); Authors thank Dr. Christine Collins for help in some of the mapping experiments.

## REFERENCES

Apkarian, A. V. and Hodge, C. J. (1989). Primate spinothalamic pathways: I. A quantitative study of the cells of origin of the spinothalamic pathway. J Comp Neurol, 288(3), 447–473. doi:10.1002/cne.902880307

Blakemore, C. and Van Sluyters, R. C. (1974). Reversal of the physiological effects of monocular deprivation in kittens: further evidence for a sensitive period. J Physiol, 237(1), 195–216. doi:10.1113/jphysiol.1974.sp010478

Boivie, J. (1979). An anatomical reinvestigation of the termination of the spinothalamic tract in the monkey. J Comp Neurol, 186(3), 343–369. doi:10.1002/cne.901860304

Churchill, J. D., Arnold, L. L. and Garraghty, P. E. (2001). Somatotopic reorganization in the brainstem and thalamus following peripheral nerve injury in adult primates. Brain Res, 910(1-2), 142–152. doi:10.1016/s0006-8993(01)02703-2

Craig, A. D. (2008). Retrograde analyses of spinothalamic projections in the macaque monkey: input to the ventral lateral nucleus. J Comp Neurol, 508(2), 315–328. doi:10.1002/cne.21672

Darian-Smith, C., Darian-Smith, I., and Cheema, S. S. (1990a). Thalamic projections to sensorimotor cortex in the macaque monkey: use of multiple retrograde fluorescent tracers. J Comp Neurol, 299(1), 17–46. doi:10.1002/cne.902990103

Darian-Smith, C., Darian-Smith, I., and Cheema, S. S. (1990b). Thalamic projections to sensorimotor cortex in the newborn macaque. J Comp Neurol, 299(1), 47–63. doi:10.1002/cne.902990104

Davidson, S., Truong, H., and Giesler, G. J., Jr. (2010). Quantitative analysis of spinothalamic tract neurons in adult and developing mouse. J Comp Neurol, 518(16), 3193–3204. doi:10.1002/cne.22392

Dutta, A., Kambi, N., Raghunathan, P., Khushu, S., and Jain, N. (2014). Large-scale reorganization of the somatosensory cortex of adult macaque monkeys revealed by fMRI. Brain Struct Funct, 219(4), 1305–1320. doi:10.1007/s00429-013-0569-8

Felleman, D. J., Nelson, R. J., Sur, M., and Kaas, J. H. (1983). Representations of the body surface in areas 3b and 1 of postcentral parietal cortex of Cebus monkeys. Brain Res, 268(1), 15–26. doi:10.1016/0006-8993(83)90386-4

Fitzgerald, M. and Jennings, E. (1999). The postnatal development of spinal sensory processing. Proc Natl Acad Sci U S A, 96(14), 7719–7722. doi:10.1073/pnas.96.14.7719

Florence, S. L., Jain, N., and Kaas, J. H. (1997). Plasticity of somatosensory cortex in primates. Seminars Neurosci., 9, 3–12.

Florence, S. L., Jain, N., Pospichal, M. W., Beck, P. D., Sly, D. L., and Kaas, J. H. (1996). Central reorganization of sensory pathways following peripheral nerve regeneration in fetal monkeys. Nature, 381(6577), 69–71. doi:10.1038/381069a0

Florence, S. L., Wall, J. T., and Kaas, J. H. (1988). The somatotopic pattern of afferent projections from the digits to the spinal cord and cuneate nucleus in macaque monkeys. Brain Res, 452(1-2), 388–392. doi:10.1016/0006-8993(88)90045-5

Galea, M. P. and Darian-Smith, I. (1995). Postnatal maturation of the direct corticospinal projections in the macaque monkey. Cereb Cortex, 5(6), 518–540. doi:10.1093/cercor/5.6.518

Garraghty, P. E. and Kaas, J. H. (1991). Functional reorganization in adult monkey thalamus after peripheral nerve injury. NeuroReport, 2(12), 747–750. doi:10.1097/00001756-199112000-00004

Granmo, M., Petersson, P., and Schouenborg, J. (2008). Action-based body maps in the spinal cord emerge from a transitory floating organization. J Neurosci, 28(21), 5494–5503. doi:10.1523/JNEUROSCI.0651-08.2008

Halder, P., Kambi, N., Chand, P., and Jain, N. (2018). Altered Expression of Reorganized Inputs as They Ascend From the Cuneate Nucleus to Cortical Area 3b in Monkeys With Long-Term Spinal Cord Injuries. Cereb Cortex, 28(11), 3922–3938. doi:10.1093/cercor/bhx256

Hubel, D. H., Wiesel, T. N., and LeVay, S. (1977). Plasticity of ocular dominance columns in monkey striate cortex. Philos Trans R Soc Lond B Biol Sci, 278(961), 377–409. doi:10.1098/rstb.1977.0050

Jain, N., Catania, K. C., and Kaas, J. H. (1997). Deactivation and reactivation of somatosensory cortex after dorsal spinal cord injury. Nature, 386(6624), 495–498. doi:10.1038/386495a0

Jain, N., Florence, S. L., and Kaas, J. H. (1995). GAP-43 expression in the medulla of macaque monkeys: changes during postnatal development and the effects of early median nerve repair. Brain research. Developmental brain research, 90(1-2), 24–34. doi:10.1016/0165-3806(96)83483-3

Jain, N., Florence, S. L., Qi, H.-X., and Kaas, J. H. (2000). Growth of new brain stem connections in adult monkeys with massive sensory loss. Proc Natl Acad Sci USA, 97, 5546–5550.

Jain, N., Qi, H., A., R., Lyon, D. C., and J.H., K. (2025a). Reorganized cortical somatotopy reflects thalamic somatotopy in monkeys with dorsal spinal cord injuries sustained early in development. Paper presented at the Soc for Neurosci, San Diego.

Jain, N., Qi, H. X., Collins, C. E., and Kaas, J. H. (2008). Large-scale reorganization in the somatosensory cortex and thalamus after sensory loss in macaque monkeys. J Neurosci, 28(43), 11042–11060. doi:10.1523/JNEUROSCI.2334-08.2008

Jain, N., Qi, H. X., Raman, A., Lyon, D., and Kaas, J. H. (2025b). Cortical reorganization following dorsal spinal injuries in newborn monkeys reveals a critical period in the development of the somatosensory cortex. Proc Natl Acad Sci U S A, 122(4), e2417417122. doi:10.1073/pnas.2417417122

Jones, E. G., and Friedman, D. P. (1982). Projection pattern of functional components of thalamic ventrobasal complex on monkey somatosensory cortex. J Neurophysiol, 48(2), 521–544. doi:10.1152/jn.1982.48.2.521

Kaas, J. H., Florence, S. L., and Jain, N. (1999). Subcortical contributions to massive cortical reorganzations. Neuron, 22, 657–660.

Kaas, J. H., Krubitzer, L. A., Chino, Y. M., Langston, A. L., Polley, E. H., and Blair, N. (1990). Reorganisation of the retinotopic cortical maps in adult mammals after lesions of the retina. Science, 248, 229–231.

Kaas, J. H., Nelson, R. J., Sur, M., Dykes, R. W., and Merzenich, M. M. (1984). The somatotopic organization of the ventroposterior thalamus of the squirrel monkey, *Saimiri sciureus*. J Comp Neurol, 226(1), 111–140. doi:10.1002/cne.902260109

Kambi, N., Halder, P., Rajan, R., Arora, V., Chand, P., Arora, M., and Jain, N. (2014). Large-scale reorganization of the somatosensory cortex following spinal cord injuries is due to brainstem plasticity. Nat Commun, 5, 3602. doi:10.1038/ncomms4602

Kutcher, M. R., and Duffy, K. R. (2007). Cytoskeleton alteration correlates with gross structural plasticity in the cat lateral geniculate nucleus. Vis Neurosci, 24(6), 775–785. doi:10.1017/S095252380707068X

LeVay, S., Wiesel, T. N., and Hubel, D. H. (1980). The development of ocular dominance columns in normal and visually deprived monkeys. J Comp Neurol, 191(1), 1–51. doi:10.1002/cne.901910102

Merzenich, M. M., Kaas, J. H., Wall, J., Nelson, R. J., Sur, M., and Felleman, D. (1983). Topographic reorganization of somatosensory cortical areas 3b and 1 in adult monkeys following restricted deafferentation. Neurosci., 8(1), 33–55. doi:10.1016/0306-4522(83)90024-6

Mountcastle, V. B., and Henneman, E. (1952). The representation of tactile sensibility in the thalamus of the monkey. J Comp Neurol, 97(3), 409–439. doi:10.1002/cne.900970302

Nelson, R. J.. and Kaas, J. H. (1981). Connections of the ventroposterior nucleus of the thalamus with the body surface representations in cortical areas 3b and 1 of the cynomolgus macaque, (Macaca fascicularis). J Comp Neurol, 199(1), 29–64. doi:10.1002/cne.901990104

Pouchelon, G., Gambino, F., Bellone, C., Telley, L., Vitali, I., Luscher, C., Holtmaat, A., and Jabaudon, D. (2014). Modality-specific thalamocortical inputs instruct the identity of postsynaptic L4 neurons. Nature, 511(7510), 471–474. doi:10.1038/nature13390

Qi, H. X., Jain, N., Collins, C. E., Lyon, D. C., and Kaas, J. H. (2010). Functional organization of motor cortex of adult macaque monkeys is altered by sensory loss in infancy. Proc Natl Acad Sci U S A, 107(7), 3192–3197. doi:10.1073/pnas.0914962107

Schwaber, M. K., Garraghty, P. E., and Kaas, J. H. (1993). Neuroplasticity of the adult primate auditory cortex following cochlear hearing loss. Am J Otol, 14(3), 252–258.

Sesma, M. A., Irvin, G. E., Kuyk, T. K., Norton, T. T., and Casagrande, V. A. (1984). Effects of monocular deprivation on the lateral geniculate nucleus in a primate. Proc Natl Acad Sci U S A, 81(7), 2255–2259. doi:10.1073/pnas.81.7.2255

Shi, T., and Apkarian, A. V. (1995). Morphology of thalamocortical neurons projecting to the primary somatosensory cortex and their relationship to spinothalamic terminals in the squirrel monkey. J Comp Neurol, 361(1), 1–24.

Simi, A., and Studer, M. (2018). Developmental genetic programs and activity-dependent mechanisms instruct neocortical area mapping. Curr Opin Neurobiol, 53, 96–102. doi:10.1016/j.conb.2018.06.007

Sloper, J. J., Headon, M. P., and Powell, T. P. (1988). A comparison of cell size changes in central and pericentral representations within the primate lateral geniculate nucleus following early monocular deprivation. Brain Res, 468(1), 61–64. doi:10.1016/0165-3806(88)90007-7

Takahata, T., Patel, N. B., Balaram, P., Chino, Y. M., and Kaas, J. H. (2018). Long-term histological changes in the macaque primary visual cortex and the lateral geniculate nucleus after monocular deprivation produced by early restricted retinal lesions and diffuser induced form deprivation. J Comp Neurol, 526(18), 2955–2972. doi:10.1002/cne.24494

Tandon, S., Kambi, N., Lazar, L., Mohammed, H., and Jain, N. (2009). Large-scale expansion of the face representation in somatosensory areas of the lateral sulcus after spinal cord injuries in monkeys. J Neurosci, 29(38), 12009–12019. doi:10.1523/JNEUROSCI.2118-09.2009

Willis, W. D., Kenshalo, D. R., Jr., and Leonard, R. B. (1979). The cells of origin of the primate spinothalamic tract. J Comp Neurol, 188(4), 543–573. doi:10.1002/cne.901880404

Wong-Riley, M. (1979). Changes in the visual system of monocularly sutured or enucleated cats demonstrable with cytochrome oxidase histochemistry. Brain Res, 171(1), 11–28. doi:10.1016/0006-8993(79)90728-5

